# Sodium ion-assisted structural lipidomics for sphingolipid profiling

**DOI:** 10.64898/2026.02.02.703417

**Authors:** Hiroaki Takeda, Daiki Asakawa, Manami Takeuchi, Hiroshi Tsugawa

**Author notes:** **Corresponding Authors** Hiroaki Takeda (H. Takeda), Hiroshi Tsugawa (H. Tsugawa). Hiroaki Takeda and Daiki Asakawa contributed equally to this work.

## Abstract

Sphingolipids are diverse lipids with sphingobases and *N*-acyl fatty acids as the hydrophobic moieties. While the importance of the in-depth elucidation of hydrophobic structures is widely recognized in lipid biology, mass spectrometry-based annotation of ceramides in the commonly used protonated form is often hindered by in-source dehydration during electrospray ionization in the heated state and variable water losses in the product ion spectrum. In this study, we investigated the sodium ion form and its product ions in ceramides with the use of electron-activated dissociation tandem mass spectrometry (EAD MS/MS) in addition to collision-induced dissociation to facilitate indepth structural elucidation. While dehydrated ions from the protonated form were frequently observed, the sodium adduct ions remained stable because of their higher activation energy compared with the protonated form, which was validated using quantum chemical calculations. Using the three adduct forms under optimized conditions increased confidence in annotating the ceramide peaks through retention-time matching. Furthermore, EAD MS/MS of the sodium adduct ions facilitated the positional determination of double bonds and hydroxyl groups in the ceramide hydrophobic moiety. Our approach is showcased by the annotation of phytoceramides with *N*-acyl 2- and 3-hydroxyl groups in mouse feces and ceramides with *N*-acyl *n*-6 very long-chain polyunsaturated 2-hydroxy fatty acids in mouse testis.

## Introduction

Lipids play essential roles in biological systems by functioning as cell membrane components, signaling molecules, and energy storage^1−3^. Their immense diversity arises from various structural categories such as phospholipids, sphingolipids, and neutral lipids, as well as various fatty acyl chains. Among them, sphingolipids are not only structural components of cell membranes but also critical mediators of cellular signaling, such as apoptosis and stress responses^4^. Dysregulation of sphingolipid metabolism has been associated with various diseases, including neurodegenerative disorders^5^, cancer^6^, atherosclerosis^7^, and metabolic diseases^8^. Although over 50,000 lipids are cataloged in the LIPID MAPS Structure Database, sphingolipids remain a particularly challenging group in analytical chemistry because of their structural complexity and functional diversity^9^. Hereafter, the lipid structure description in this study follows the guidelines of the Lipidomics Standards Initiative^10^, and the scope for the acyl chain description is to determine the carbon number, degree of unsaturation, and carbon-carbon double bond (C=C) and hydroxy (OH) positions.

Mass spectrometry (MS)-based metabolome atlases have become widely available online, providing valuable insights into the spatial distribution of metabolites and accelerating biological research^11−13^. As demonstrated by the MetaboAtlas21 database, ceramides (Cers) exhibit distinct localization patterns across various organs, tissues, blood, and feces depending on their acyl chain compositions^14^. For example, Cer 18:1;O2/18:0, known as the NS type, is predominantly distributed in the brain, whereas Cer 18:1;O2/16:0 shows a markedly different distribution across the kidney, stomach, pancreas, intestine, and testis. Additionally, OH group-rich Cers tend to accumulate in the skin, gut, and feces, suggesting a role in intestinal metabolism and microbial interactions^15,16^. While the *N*-acyl chains of Cers are primarily composed of saturated and monounsaturated fatty acids (FAs), polyunsaturated FA-containing Cers are notably enriched in the testis, indicating potential tissue-specific functions^17^. Given this structural diversity, a comprehensive sphingolipid analysis is essential to elucidate their biological roles, uncover their regulatory mechanisms, and explore potential therapeutic targets for various physiological and pathological conditions.

Sphingolipids are often analyzed after alkaline hydrolysis of glycerolipids^18^, which exploits the higher stability of amide bonds than ester bonds. However, their structural diversity makes selective enrichment a challenge. To address this issue, recent studies have introduced titania (TiO_2_)-based methods, such as TiO_2_-coated slides for liquid microjunction surface sampling^19^, TiO_2_- and zirconia (ZrO_2_)-coated solid-phase extraction (SPE) silica column^20^, and magnetic TiO_2_ nanoparticle-based enrichment^21^. TiO_2_ enables the enrichment of sphingolipids and the removal of phosphorylated compounds via Lewis acid-base interactions^22^. These SPE techniques improve the detection of low-amount species and facilitate structural characterization of sphingolipids, including C=C and OH positions, through Paternò-Büchi reaction or epoxidation^23,24^ and different modalities of mass fragmentation strategies^25^.

Lipid fragmentation using tandem MS (MS/MS) can be categorized into two types. The first is partial fragmentation, which generates product ions specific to features such as double bonds and fatty acyl chains. Ultraviolet photodissociation^26^, ozone-induced dissociation^27^, and oxygen attachment dissociation (OAD)^28^ are frequently used in in-depth lipidomics studies, with ozone-induced dissociation and OAD being particularly effective for generating C=C position-specific fragment ions. The other is full fragmentation, in which most chemical bonds, including carbon-carbon single bonds (C–C), can be cleaved, which is facilitated by hydrogen abstraction dissociation^28^ and electron-activated dissociation (EAD)^29−33^. Recently, we developed a dual collision-induced dissociation (CID) and OAD method, termed OAciD, in which atomic oxygen and hydroxyl radicals selectively oxidize double bonds, whereas residual water vapor promotes CID, enabling detailed analysis of lipid subclasses, fatty acyl compositions, and C=C positions with a limited set of diagnostic fragments^34^. OAciD improves in-depth lipidomics even for incompletely separated isomers. In contrast, EAD facilitates homolytic cleavage, enabling the assignment of C=C and OH positions as well as acyl arrangements. Recent improvements in electron-beam optics and trap-and-release strategies have significantly increased the sensitivity of quadrupole time-of-flight MS (QTOFMS)^35^. We further formulated EAD MS/MS spectral patterns for various lipid subclasses, whose algorithms were integrated into MS-DIAL 5 for automated structural annotation^36^.

The EAD MS/MS technique is limited to several lipid subclasses in reverse-phase liquid chromatography (RPLC)-based untargeted lipidomics. Since different isomers, including those with varying FA compositions, C=C positions, and *sn*-positions, often co-elute within the same retention-time ranges, the interpretation of the product ion spectra becomes difficult because (1) most bonds can be cleaved, (2) each bond cleavage can generate three types of fragment ions, including hydrogen (H)-loss fragments, radicals, and H-gain fragments, and (3) homolytic cleavage-derived fragment ions are also generated from the polar head loss form of precursor ions in complex lipids, resulting in the generation of non-intuitive spectra if multiple isomers are co-eluted. Currently, several lipid subclasses with strong charge stabilization of the quaternary ammonium cations, such as phosphatidylcholine and sphingomyelin (SM), are suitable targets for EAD MS/MS-based structural elucidation^36^. Furthermore, the C=C and OH positions of FA metabolites were determined after derivatization with *N,N*-dimethylethylenediamine, which readily forms a cation through structural change^37^.

In sphingolipid analysis, RPLC/MS characterizes 30 types of free and protein-bound Cers from human and mouse stratum corneum^15^. Cers often undergo dehydration during electrospray ionization (ESI) or fragmentation due to the presence of OH groups. These undesired structural modifications can lead to misannotation, reduced sensitivity of the precursor ions, loss of structure-specific fragments, and increased complexity of the structural analysis. During ESI, stable ionized Cers can be generated using a lithium-containing solvent, which improves the structural analysis with higher collision energy in linear ion trap MS^38^. EAD MS/MS of protonated Cers also involves a combination of vibrational activation cleavage (as seen in CID), charge-remote fragmentation initiated by electronic activation (as observed in EAD), and dehydration processes occurring during fragmentation. Sodium adducts offer a valuable alternative for simplifying the mass fragmentation of sphingolipids by providing different adduct sites and dissociation rates for dehydration. This not only simplifies the interpretation of Cer structures but also clarifies the existence of lipid isomers in the MS data.

In this study, we developed an analytical workflow for the in-depth profiling of sphingolipids that uses sodium adduct molecules, in addition to proton adduct molecules, to effectively perform peak annotation and provide deeper structural insights. Quantum chemistry calculations based on density functional theory (DFT) were employed to elucidate the fragmentation pathways and dissociation rates of the target ions. Furthermore, we used the sodium adduct EAD to facilitate in-depth structural elucidation of OH group-rich Cers composed of phytosphingosine and *N*-acyl hydroxy FAs, in addition to very long-chain polyunsaturated FAs (VLC-PUFAs) containing Cers. Its utility was demonstrated through lipidomics studies of the mouse testis and feces.

### Experimental Section

#### Chemicals and reagents

Reagent-grade 25.0% ammonia solution (NH_3_) and chloroform (CHCl_3_), 1 mol/L ammonium acetate solution for high-performance LC, formic acid (HCOOH) for LC/MS, acetonitrile (MeCN), methanol (MeOH), 2-propanol (IPA), and ultrapure water (H_2_O) for QTOFMS were obtained from FUJIFILM Wako Pure Chemical Corp. (Osaka, Japan). TiO_2_ and ZrO_2_-coated monolithic silica MonoSpin Phospholipid (<800 μL; through pore, 5 μm; mesopore, 10 nm; φ 4.2 mm × 1.5 mm) was purchased from GL Sciences, Inc. (Tokyo, Japan) for sphingolipids enrichment. Synthetic standards were purchased from Avanti Polar Lipids, Inc. (Alabaster, AL, USA).

#### Animals

The same animals from the previous study were used in this study^20^. All animal experiments were performed in accordance with an ethical protocol approved by the Tokyo University of Agriculture and Technology (R5-50). C57BL/6J male mice were purchased from SLC (Shizuoka, Japan). The mice were fed CE-2 chow (CLEA Japan, Tokyo, Japan). The testis and feces were harvested, immediately frozen after dissection, and stored at −80 °C until extraction.

#### Liquid extraction

Because biphasic liquid extraction (a modified Bligh and Dyer method^39^) showed lipid profiles comparable to those obtained with monophasic extraction, except for testicular triacylglycerols^20^, the present study employed a simple monophasic liquid extraction as the SPE pretreatment. Lyophilized samples (approximately 15.2 mg of feces and 9.6 mg of testis) were mixed with 200 μL of MeOH and extracted by ultrasonication for 5 min. One hundred microliter of CHCl_3_ was added, and the solution was vortexed for 1 min and extracted on ice for 60 min. The solution was further mixed with 20 μL of H_2_O and vortexed for 1 min. The supernatant was collected after centrifugation at 16,000 × *g* for 5 min at 4 °C. The lipid extracts were dried using a centrifuge evaporator.

#### Solid phase enrichment

The SPE method was developed using the MonoSpin Phospholipid column^20^. A dried lipid extract was dissolved by adding 100 μL of CHCl_3_. The SPE columns were conditioned with 200 μL of CHCl_3_ before loading the lipid extract. Fifty microliters of the lipid extract dissolved in CHCl_3_ were applied to the SPE columns and centrifuged at 500 × *g* for 2 min at 10 °C to elute neutral lipids such as triacylglycerols. To remove the neutral lipids, 200 μL of CHCl_3_ was added to the SPE columns and centrifuged under the same conditions. The SPE columns were then loaded with 200 μL MeOH/HCOOH (99:1, v/v) and centrifuged at 500 × *g* for 5 min at 10 °C to elute sphingolipids, except for SMs. The fraction was collected twice to increase recovery. Acidic MeOH fractions collected by SPE were dried in a centrifugal evaporator and reconstituted in 60 µL of MeOH for LC-MS analysis.

#### MS/MS acquisition for synthetic compounds

The LC/QTOFMS system comprised an Exion LC and a ZenoTOF 7600 with an ESI ion source (SCIEX, Framingham, MA, USA). Previously developed high-throughput LC gradient conditions were used to acquire the MS/MS spectra of the synthetic standards^40^. The LC conditions were as follows: injection volume, 1 μL; mobile phase, MeCN/MeOH/H_2_O (1:1:3, v/v/v) (A) and MeCN/IPA (1:9, v/v) (B) (both contained 10 nM of ethylenediaminetetraacetic acid and 5 mM of ammonium acetate); flow rate, 600 μL/min; column; Unison UK-C18 MF (50 × 2.0 mm, 3 μm, Imtakt Corp., Kyoto, Japan); gradient, 0.1% (B) (0.1 min), 0.1–15% (B) (0.1 min), 15–30% (B) (0.9 min), 30–48% (B) (0.3 min), 48–82% (B) (4.2 min), 82–99.9% (B) (1.3 min), 99.9% (B) (0.2 min), 99.9–0.1% (B) (0.1 min), 0.1% (B) (1.4 min); column oven temperature, 65 °C. The column eluent was mixed with 50 μL/min of 50% methanol containing sodium acetate (40 μM) post-column when forming the sodium adduct ion. MS conditions were applied as previously described^40^. Source and gas parameters were as follows: ion source gas 1, 40 psi; ion source gas 2, 80 psi; curtain gas, 30 psi; CAD gas, 7; source temperature, 250 °C; spray voltage, 5500 V. MRMHR were as follows: Q1 resolution, unit; electron-beam current, 7000 nA; Zeno threshold, 1,000,000 cps; accumulation time, 100 ms; declustering potential, 80 V; collision energy, 40 ± 15 V for CID and electron kinetic energy, 8, 10, 12, 14, 16, 18, and 20 eV for EAD; EAD-RF, 150 Da; reaction time, 98 ms

#### Sphingolipid profiling

The LC conditions were as follows: injection volume, 2 μL; mobile phase, MeCN/MeOH/H_2_O (1:1:3, v/v/v) (A) and MeCN/IPA (1:9, v/v) (B) (both containing 5 mM ammonium acetate and 10 nM ethylenediaminetetraacetic acid); flow rate, 300 μL/min; column, Unison UK-C18 MF (50 mm × 2.0 mm, 3 μm, Imtakt Corp., Kyoto, Japan); gradient, 0.5% (B) (1 min), 0.5–40% (B) (4 min), 40–64% (B) (2.5 min), 64–71% (B) (4.5 min), 71–82.5% (B) (0.5 min), 82.5–85% (B) (6.5 min), 85–99% (B) (1 min), 99% (B) (2 min), 99–0.5% (B) (0.1 min), 0.5% (B) (2.9 min); column oven temperature, 45 °C. The column eluent was mixed with 10 μL/min of 50% methanol containing 200 μM sodium acetate post-column when forming the sodium adduct ion. The source and gas parameters for the MS conditions were the same as those used for the MS/MS acquisition. TOF-MS for both CID and EAD were as follows: mass range, 70–1250 Da; declustering potential, 80 V; collision energy, 10 V; accumulation time, 200 ms. TOF-MS/MS for CID was as follows: Q1 resolution, unit; mass range, 70–1250 Da; declustering potential, 80 V; collision energy, 40 ± 15 V; accumulation time, 50 ms; Zeno threshold, 2,000,000 cps. TOF-MS/MS for EAD was as follows: Q1 resolution, unit; mass range, 100–1250 Da; declustering potential, 80 V; collision energy, 10 V; accumulation time, 100 ms; kinetic energy, 10, 14, or 18 eV; electron-beam current, 7000 nA; Zeno threshold, 1,000,000 cps; EAD-RF, 150 Da; reaction time, 98 ms.

#### Data analysis

Peak picking of lipids was performed using MS-DIAL version 5.1^36^. The data processing parameters and adduct ions used to calculate the lipid amounts are listed in Table S1. The annotated results were verified using the retention times and MS/MS spectra of the peaks in positive-ion mode. The annotation results are provided in Tables S2 and S3.

#### Lipid nomenclature

The nomenclature used in this study is described in accordance with the Lipidomics Standards Initiative^10^. Cer types were defined based on the structure of the long-chain base (dihydrosphingosine (SPB 18:0;1OH,3OH), sphingosine (SPB 18:1;1OH,3OH), and phytosphingosine (SPB 18:0;1OH,3OH,4OH)) and *N*-acyl FA (non-hydroxy FA, *alpha*-hydroxy (2-OH) FA, and *beta*-hydroxy (3-OH) FA). By combining these structures, Cer-NS (non-hydroxy FA and sphingosine), Cer-NP (non-hydroxy FA and phytosphingosine), Cer-AS (*alpha*-hydroxy FA and sphingosine), Cer-AP (*alpha*-hydroxy FA and phytosphingosine), and Cer-BP (*beta*-hydroxy FA and phytosphingosine) are described according to the following rules. For example, Cer 18:1;O2/16:0 indicated that individual fatty acyl chains were determined using MS/MS. The detailed position is described as Cer 18:1(Δ4);1OH,3OH/16:0 when fragment ions derived from the C=C and OH positions are obtained using EAD.

#### Quantum chemistry calculation

All electronic structures of the lipids and their fragments were calculated using the Gaussian 16 suite^41^. First, the optimized geometries of the proton and sodium adducts Cer 18:1(4E);1OH,3OH/16:0 were determined using DFT with the ωB97X-D hybrid functional^42^ and 6-31+G(d,p) basis set. To obtain the transition-state conformation for dehydration, the C–O bond length was gradually increased until an energy maximum was reached, after which the obtained conformation was further optimized at the ωB97X-D/6-31+G(d,p) level of theory to find the saddle point. Frequency analysis of the transition state geometries was conducted to confirm the presence of an imaginary vibration frequency. To determine the energetics required for fragmentation, the relationships among the transition states, reactants, and intermediates were verified using intrinsic reaction coordinate analysis, starting from the transition-state geometry. The frequency analysis results were also used for zero-point energy corrections. Microcanonical rate constants were computed based on the Rice−Ramsperger−Kassel−Marcus theory using the MassKinetics program version 1.21^43^. The homolytic allyl cleavage and N–C bond cleavage of the sodium adduct Cer 18:1(4E);1OH,3OH/16:0 was calculated using spin-flip DFT at the ωB97X-D/6-31+G(d,p) theoretical level. To obtain the potential energy surface in the ground state, the corresponding bond length was increased incrementally in 0.1 Å steps. The potential energy surfaces of the excited singlet states were calculated using time-dependent DFT (TD-DFT) with ωB97X-D/6-31+G(d,p) level of theory, using conformations on ground states obtained using spin-flip DFT.

## Results and Discussion

### Analytical workflow for in-depth sphingolipid profiling

We employed SPE-based lipid enrichment and EAD MS/MS for sphingolipid profiling (Figure 1). Our study aimed to elucidate the hydrophobic structures of Cers from the product ion spectra (Figure 1a). We have previously developed a practical SPE protocol for the fractionation of neutral lipids and steryl esters (with nonpolar functional groups), sphingolipids and FA metabolites (with hydroxyl or carboxyl groups), and phospholipids (with phosphate groups) using a TiO_2_- and ZrO_2_-coated SPE silica column (Figure 1b)^20^. This SPE column enables a two-dimensional separation mechanism, where silica provides electrostatic interactions and metal oxides facilitate coordination interactions. When the lipid extracts dissolved in CHCl_3_ are applied to the column, hydrophobic glycerolipids and steryl esters are excluded because of their low dielectric constants. In contrast, sphingolipids (excluding SM) and FA metabolites are efficiently eluted with MeOH containing volatile HCOOH, whose carboxyl group competes for interactions with the stationary phase, whereas phospholipids remain retained. Throughput and purification costs are reduced by eliminating the removal step for counter-substances, as HCOOH can be removed by drying.

**Figure 1.**
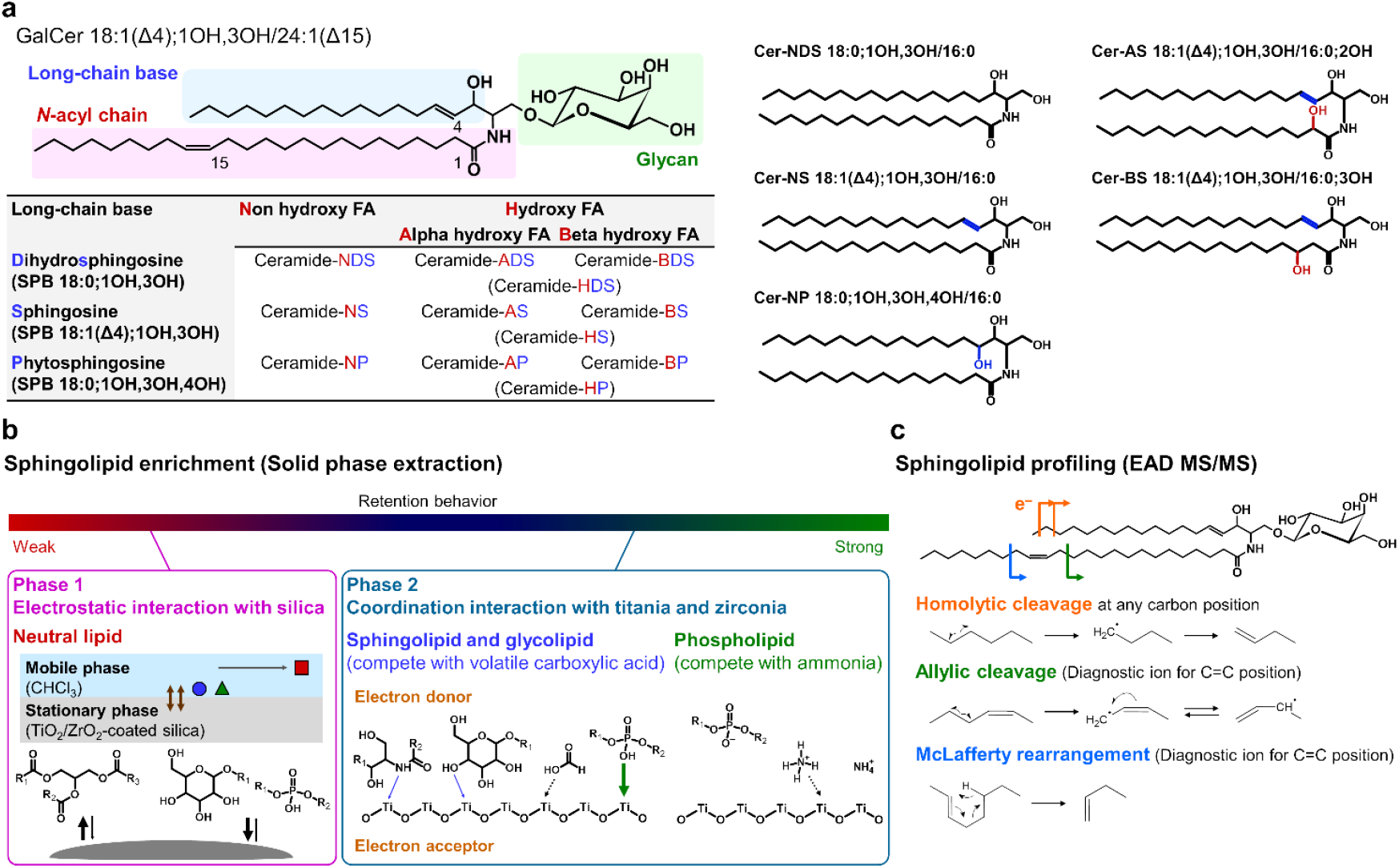
Analytical workflow for in-depth sphingolipid profiling by SPE and EAD MS/MS. (**a**) Structure of Cer types detected in mouse testis and feces extracts. Nomenclature of these Cer types is determined based on the initial alphabet of the long-chain base and the *N*-acyl chain. (**b**) Sphingolipid enrichment using TiO_2_ and ZrO_2_-coated monolithic silica columns. (**c**) EAD MS/MS fragmentation for sphingolipid structural analysis.

EAD cleaves C–C covalent bonds in fatty acyl chains via homolytic cleavage at kinetic energies of 8–20 eV^33^. This process produces radical fragments at 14.02 Da intervals, along with H-gain and H-loss fragments at each covalent bond position. At the C=C positions, the formation of allyl radical fragments, stabilized by delocalized electrons, and H-loss fragments generated through the McLafferty rearrangement (involving *γ*-position hydrogen transfer) is significantly enhanced (Figure 1c), showing a three-to five-fold increase in abundance compared with C–C positions. These characteristics enable detailed structural elucidation by interpreting the full MS/MS spectra, despite the relatively low fragmentation efficiency (approximately 1–5% of the residual precursor ion).

### Elucidation of the dehydration pathway for proton and sodium adduct ion forms in Cer-NS

Because the EAD MS/MS spectrum of protonated Cers shows fragment ions originating from both electronic and vibrational activation, spectral interpretation is particularly challenging. We first investigated the relationship between the kinetic energy and the observed fragment ions, and found that the highest number of fragments from the protonated and sodiated ion forms of Cer 18:1(Δ4);1OH,3OH/16:0 was observed at 14 eV (Figure 2a). Both the total number of fragments and their intensities decrease at energies above 16 eV. In terms of fragment ion production, the optimal kinetic energy (14 eV) was consistent with our previous findings, mostly for phospholipids and neutral lipids^36^. The EAD of the protonated precursor undergoes dehydration to produce [M+H−H_2_O]^+^. In addition, the fragment ions derived from long-chain bases, including sphingobase (SPB) 18:1;O2−CH_4_O_2_ (*m/z* 252.27), SPB 18:1;O2−2H_2_O (*m/z* 264.27), SPB 18:1;O2−H_2_O (*m/z* 282.28), and palmitamide (*m/z* 256.26), are observed in the EAD MS/MS spectrum (Figure 2b). These fragment ions also appear in CID MS/MS spectra from both precursor ions of [M+H]+ and [M+H−H_2_O]+. These results indicate that the vibrational activation of [M+H]^+^ generates fragment ions at *m/z* 252.27, 264.27, and 282.28 with [M+H−H_2_O]^+^ as an intermediate.

**Figure 2.**
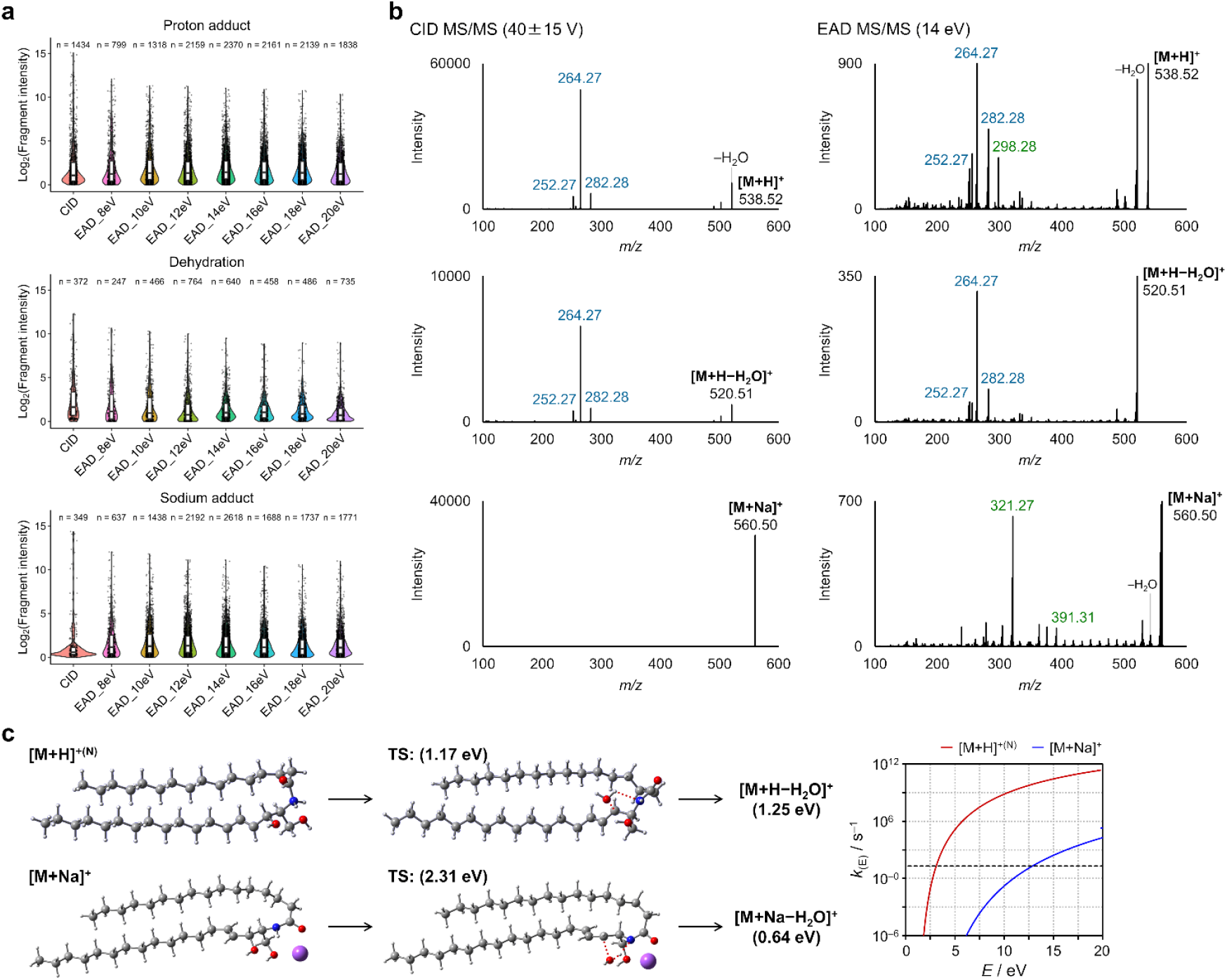
MS/MS fragmentation of Cer-NS 18:1(Δ4);1OH,3OH/16:0. (**a**) Distribution of fragment ion numbers and their intensities for proton and sodium adducts under CID and EAD conditions. Black circles indicate individual fragments with an intensity greater than 1, and the number of such fragments is shown at the top of each plot (n = xx). (**b**) EAD MS/MS of Cer-NS 18:1(Δ4);1OH,3OH/16:0 with proton adduct, its dehydrated form, and sodium adduct. Navy blue and green fragments originate from major vibrational and electronic activations, respectively. (**c**) Calculated dehydration pathway of N-protonated or sodiated Cer-NS 18:1(4E);1OH,3OH/16:0 and its *k*_(E)_ values. Carbon, hydrogen, oxygen, nitrogen, and sodium atoms are represented by the gray, white, red, blue, and purple spheres, respectively. Activation energies of transition states and enthalpy changes between reactants and products were calculated for N-protonated and sodium adduct forms.

As the initial fragmentation step of protonated Cers by vibrational activation can be dehydration, we investigated the dehydration mechanism of protonated Cers using DFT calculations. First, the global minimum geometries of the protonated Cers were searched using the ωB97X-D/6-31+G(d,p) level of theory. The calculations revealed that protonated Cer 18:1(4E);1OH,3OH/16:0 exists in two states: O-protonated (O-protomer) and N-protonated (N-protomer) forms. Because the N-protomer is 0.04 eV more stable than the O-protomer, we focused on the former (Figure S1). During dehydration, the OH group most readily released was at the Δ3 position in the protonated Cers. The TS barrier for the dehydration of N-protomers was calculated as 1.17 eV (Figure 2c). In EAD, the reaction time between precursor ions and electrons is set to be 100 ms. Therefore, the effective fragmentation timescale (τ) should be shorter than this reaction period. In this study, τ was assumed to be 50 ms. Since ion dissociation occurs as a unimolecular dissociation process, the dissociation yield of the precursor ion is exp(– *k*_(E)_τ). In this study, the appearance energy (*E*_app_) for dissociation is defined as the internal energy at which the dissociation yield of the precursor ion becomes the reciprocal of Euler’s constant, *e*, i.e., where *k*_(E)_ equals the reciprocal of τ, that is 20. The *E*_app_ of dehydration from the N-protomers was 3.1 eV (Figure 2c). Dehydration of the N-protomers occurs via heterolytic bond cleavage between carbon and 3OH, followed by binding to excess protons (Figure 2c). In addition to the N-protomers, the O-protomers also undergo dehydration (Figure S1). Therefore, the use of sodiated Cers instead of protonated Cers as precursor ions suppresses the dehydration reaction. The TS barrier and *E*_app_ value for the dehydration of sodiated Cer were obtained as 2.31 and 12.9 eV, respectively (Figure 2c). As expected, dehydration of sodiated Cers did not occur during EAD and CID because of the high *E*_app_ value. These findings suggest that the formation of sodium adducts during ESI effectively suppresses undesirable dehydration, thereby enhancing the accuracy of peak annotation and structural analysis.

### Homolytic cleavage of Cer-NS for structural characterization

EAD can induce homolytic bond cleavage, and the resulting fragment ions can reveal the position of the OH group. The fragment ion at *m/z* 298.27 indicates an OH group at the Δ3 position in the long-chain base of Cer and would be produced by homolytic cleavage of the C–C bond (Figures 2b). Although fragment ions generated through homolytic cleavage, which are informative for elucidating Cer structures, were detected in the EAD MS/MS spectrum, these diagnostically valuable ions were obscured by intense fragments arising from vibrational activation. This vibrational activation is often induced by mobile protons in protonated adducts. In addition, homolytic cleavages from both precursor ions and their dehydrated forms were observed simultaneously in the protonated precursors, further complicating the precise determination of the C=C position.

In contrast, the CID of sodiated Cers did not provide any fragment ions owing to the lack of mobile protons. The conversion of protons into sodium in the precursor ion strongly suppressed the fragmentation induced by vibrational activation, even during the EAD process. As a result, fragment ions useful for elucidating Cer structures were selectively observed in the EAD MS/MS spectrum of the sodiated Cers. (Figure 2b). The use of sodiated Cers as a precursor ion in EAD MS/MS enhances the visibility of lipid structural features, including C=C and OH positions, compared with protonated adducts, their dehydrated forms, and acetate adducts, which are used for the effective structural elucidation of Cers in CID-based lipidomics studies (Figures 2b and S2)^44^. The sensitivity of the homolysis-derived fragment ions at each covalent bond increased upon the application of sodium adduct ions, although the ionization efficiency at the MS1 peak remained constant between the proton and sodium adduct ions (Figures 3a, 3b, and S2b). The fragmentation efficiency at each covalent bond remained consistent, as indicated by the coefficients of variation of the radical fragments (20.4%) and H-loss fragments (14.9%) across positions n-2 to n-11 (Figure 3b). In contrast, the proton adducts showed significantly higher coefficients of variation: 102.2% for the radical fragments and 36.6% for the H-loss fragments (Figure 3b). This constant fragmentation efficiency with sodium adducts reduces the risk of misannotation by preventing masking of specific fragments derived from the C=C positions. Furthermore, the fold change in C=C-derived specific fragments, calculated as the ratio between the allyl radical fragment and the average of other radical fragments, increased from 3.3-fold (with the proton adduct ion) to 5.0-fold (with the sodium adduct ion), enabling a more precise characterization of the C=C positions (Figure 3b). Sodium adduct EAD also enabled the precise characterization of the OH group at the Δ3 position (*m/z* 321.26) and the double bond at the Δ4 position (*m/z* 391.31) in the long-chain base of Cer 18:1(Δ4);1OH,3OH/16:0 (Figure 3a). The total number of fragments was greater for the sodium adduct ion than for the proton adduct ion, despite the absence of vibrational activation-derived fragments in the sodium form (Figure 2a). In addition to reducing unnecessary fragment generation, the use of sodium adduct ions minimizes contamination from the spectral noise patterns observed with protonated adduct ions.

**Figure 3.**
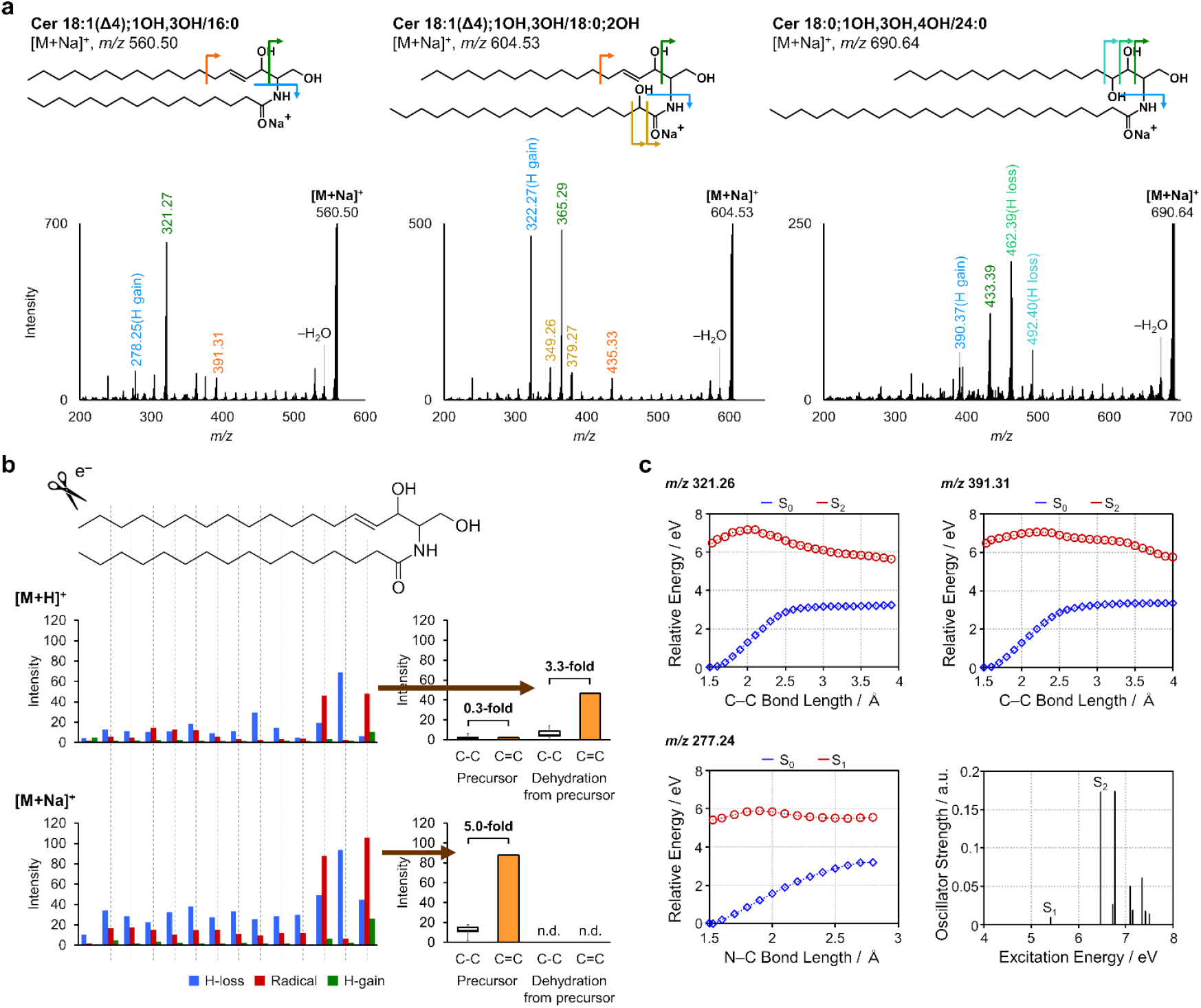
Advanced structural analysis using sodium adduct EAD MS/MS. (**a**) Sodium adduct EAD MS/MS spectra of Cers. (**b**) Intensities of three types of fragments derived from covalent bonds (radical, H-gain, and H-loss) needed to annotate the C=C position. [M+H]^+^ corresponds to homolytic cleavage products of dehydrated precursor ions, while [M+Na]^+^ corresponds to cleavage products directly from the intact precursor ions. The right panels show the intensity balance at each C–C bond and allyl radical position of the C=C bond. Both the MS/MS fragments derived from the intact precursor ions and their dehydration products were evaluated. (**c**) The potential energy surface for homolytic bond cleavage of the sodiated Cers. The ground and excited states are shown in blue and red lines, respectively. Formation of fragment ions at *m/z* 321.26, 391.31, and 277.24 was calculated. The bottom-right panel illustrates the relationship between the electronic excitation energy of sodiated Cers and its oscillator strength. The first and second excited states correspond to the π–π* excitation of amide and C=C bonds, respectively.

Next, we examined the fragment ions at *m/z* 321.26 and 391.31, which originated from allylic cleavage. Because these ions are produced by homolytic bond cleavage, the fragmentation mechanisms were investigated using spin-flip DFT. Assuming dissociation occurs at an interatomic length where the energy gradient (interatomic force) becomes zero, the required energy for the formation of the fragment ions at *m/z* 321.26 and 391.31 was 3.24 and 3.35 eV, respectively (Figure 3c). According to the calculation of *k*_(E)_ for the corresponding fragmentations, *k*_(E)_ equals the reciprocal of τ, that is 20, when the ion internal energy was approximately 18.5 eV. The results indicate that the ground-state sodiated Cer requires internal energy exceeding 18.5 eV to generate fragment ions at *m/z* 321.26 and 391.31. As the kinetic energy was set at 14 eV under the experimental conditions, the allylic bond cleavage was not due to the vibrational excitation of the ground state.

We focused on the fragmentation behavior of the singlet excited states to understand the mechanism of allylic bond cleavage. It was found that the second singlet excited state (S_2_) of sodiate Cers corresponds to the π–π* excitation of the C=C bond, and was related to allylic cleavage (Figure 3c). At the optimized geometry with a bond length of 1.53 Å, the excitation energy to the S_2_ state was calculated to be 6.49 eV. The potential energy of the S_2_ state increases slightly as the allylic bond lengthens. The allylic bond leading to the ion at *m/z* 321.26 reaches a maximum of 7.16 eV at 2.0 Å, whereas the bond producing the ion at *m/z* 391.31 shows a maximum of 7.05 eV at 2.2 Å. Consequently, the barrier for the allylic cleavages of the sodiated Cer on the S_2_ states was only approximately 0.5 eV, suggesting that the π–π* excitation of the C=C bond immediately provides the fragment ion at *m/z* 321.26 and 391.31.

In addition, the fragment ion at *m/z* 277.25 originated from the homolytic cleavage of the N–C bond, and the subsequent inter-fragment proton transfer provided the fragment ion at *m/z* 278.25. The corresponding bond cleavage was related to the first singlet excited state (S_1_), which corresponds to the π–π* excitation of amide bonds. At the optimized geometry, the excitation energy to the S_1_ state was calculated to be 5.41 eV. As the N–C bond lengthens, the potential energy increases and reaches a maximum of 5.88 eV at 1.9 Å (Figure 3c). The barrier for the N–C bond cleavages of the sodiated Cer on the S_1_ states was only 0.47 eV (Figure 3c). Consequently, fragment ions at *m/z* 277.25 and 278.25 were produced in the S_1_ state.

The ion intensities of the fragment ions at *m/z* 277.25 and 278.25 were lower than those of the ions at *m/z* 321.26 and 391.31, although the excitation energy of S_1_ was lower than that of S_2_. The oscillator strength of the S_1_ state was lower than that of the S_2_ state, suggesting that S_2_ was generated more readily than S_1_ (Figure 3c). The formation of fragment ions at *m/z* 277.25, 321.26, and 391.31 in EAD MS/MS of sodiated Cers can be understood in terms of electronic activation followed by homolytic allylic cleavage.

### Investigating the sodium adduct-derived EAD MS/MS behaviors for the other Cer types

These fragmentation rules were evaluated for AS- and NP-type Cers (Cer-AS and Cer-NP, respectively) using synthetic standards (Figures 3a and S3). The degree of dehydration varies among the Cer types, primarily depending on the presence of a double bond in the long-chain base. For example, NS type Cer (Cer-NS) is highly dehydrated, whereas NDS-type Cer (Cer-NDS) and Cer-NP exhibit less dehydration^15^. Our sodium LC-MS system substantially suppressed the dehydration of Cer-NP during ESI (Figure S2b). The peak areas of the dehydration products of Cer-NS and Cer-AS were 27.1% and 49.1%, respectively, compared with the values for the proton adducts (Figure S2b). Nevertheless, OH position-related fragment ions such as *m/z* 349.26 of the *N*-acyl chain and *m/z* 365.29 of the long-chain base for Cer 18:1;1OH,3OH/18:0;2OH, and *m/z* 433.39, 462.39, and 492.40 of the long-chain base for Cer 18:0;1OH,3OH,4OH/24:0 were clearly characterized, regardless of their location on the long-chain base or *N*-acyl chain moieties (Figure 3a). In addition to providing precise structural characterization, our sodium LC-MS system mitigated the risk of misannotation caused by dehydration during ESI.

In structural analysis of Cer 18:1(Δ4);1OH,3OH/16:0, the long-chain base composition induced by vibrational activation (*m/z* 252.27, 264.27, and 282.28) and C=C and OH positions derived from electronic activation (*m/z* 391.31 and 321.26, respectively) were the key diagnostic fragments (Figure 4a). Consistent with the total number of fragments observed at each energy level (Figure 2a), the fragmentation efficiency of these structure-specific patterns was high at 10–14 eV, indicating that this energy condition achieves a balance between spectral information content and detection sensitivity (Figure 4b). This rule also applies to other Cer types. A summary of the fragmentation patterns observed for the three types of adduct ions is presented in Figure 4c. Simultaneous analysis of Cers using the protonated and sodium forms of EAD MS/MS enabled highly reliable annotation of the long-chain base compositions and delta positions of the C=C and OH groups (Figure 4c). The feasibility of using sodium adduct ions, as assessed by quantum chemical calculations, was clearly reflected in the data-driven approach to EAD-based structural analysis.

**Figure 4.**
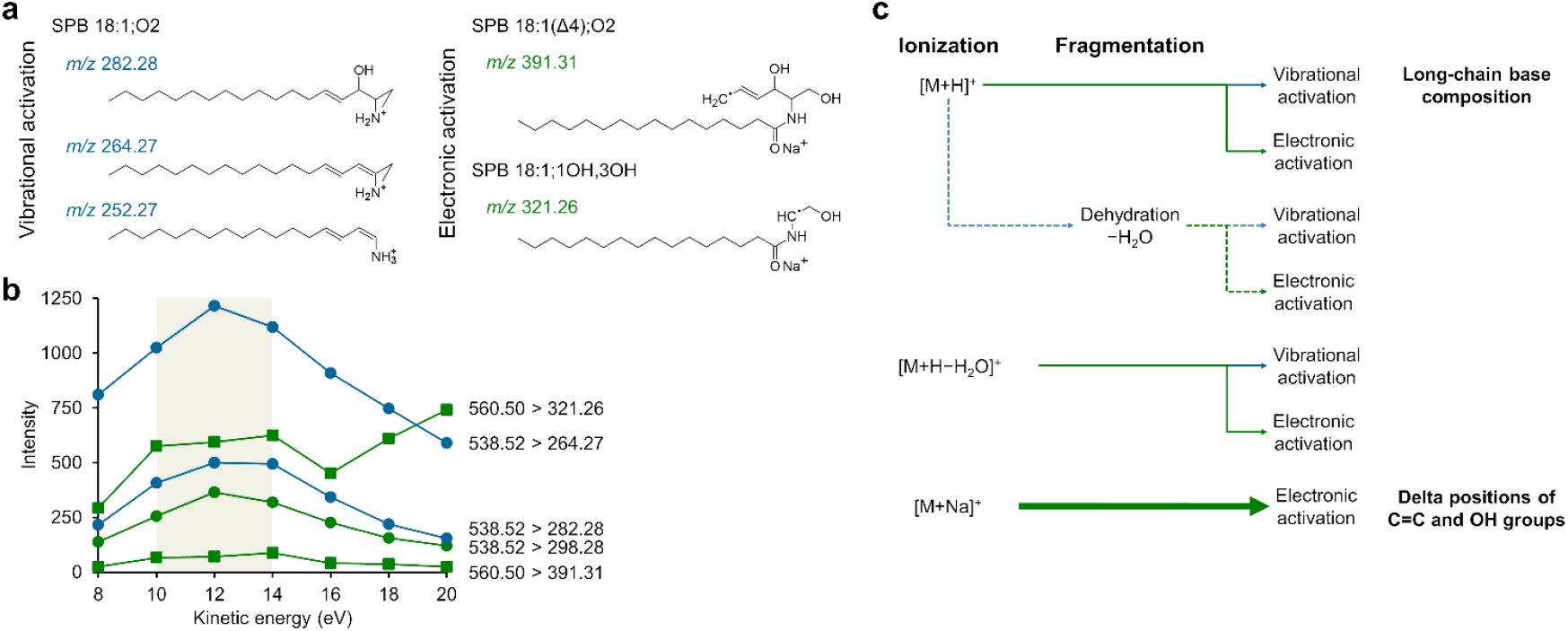
Summary of key fragment ions for structural analysis. (**a**) Key fragment ions for characterization of sphingolipid structure derived from long-chain base composition and C=C and OH positions on long-chain base and *N*-acyl chain. (**b**) Intensity behavior of key fragment ions detected from Cer-NS 18:1(Δ4);1OH,3OH/16:0 with increasing kinetic energy. Kinetic energy of 10–14 eV provides specific fragment ions with high sensitivity. (**c**) Flowchart of fragment ions detected under each adduct ion type.

### Sphingolipid profiling of biological samples using sodium adduct ions

Our analytical platform, combined with the SPE protocol and untargeted lipidomics using the MS-DIAL 5 software, enabled the detection of minor glycolipids that are often masked by more abundant lipids^20^. For instance, OH group-rich Cers with long-chain bases such as SPB 18:0;O3 and/or *N*-acyl hydroxy FAs were annotated in the sphingolipid fraction of mouse fecal extracts. However, their detailed structures were difficult to resolve using conventional CID MS/MS because of the coexistence of [M+H]^+^ and [M+H−H_2_O]^+^ ions, which complicate peak annotation. This problem was caused by the high source temperatures of the ESI instruments. The dehydration product of Cer 18:1(Δ4);1OH,3OH/24:0;2OH ([M+H−H_2_O]^+^) and the proton adduct ion of Cer 18:1(Δ4);1OH,3OH/24:1(Δ15) ([M+H]^+^) share the same exact mass (*m/z* 648.63, C_42_H_82_NO_3_). Although RPLC often resolves these peaks based on differences in the double bond count, annotation becomes challenging in complex biological samples containing numerous OH group-rich Cers. In contrast, the dehydration products of sodium adduct ions ([M+Na−H_2_O]^+^) are undetectable under conventional conditions. Leveraging this property, the composition of Cer types in biological samples can be clearly annotated using [M+H]^+^, [M+H−H_2_O]^+^, and [M+Na]^+^ ions detected at the same retention time

Our previous study demonstrated that proton adduct EAD has a limited capacity to reveal detailed structures, except for phosphatidylcholines and SMs^36^. However, detailed structural characterization was achieved using sodium adduct EAD MS/MS, as described in this study. We analyzed the sphingolipid fractions from the feces and testis samples (Tables S2 and S3). OH group-rich Cers, including Cer-NP and Cer-AS, are commonly detected in feces^11,16^. In addition to these types, sodium adduct Cer 18:0;O3/24:0;O (*m/z* 706.63) detected in the sphingolipid fraction of mouse feces enabled precise characterization of OH groups at the Δ3 and Δ4 positions (*m/z* 478.39 and 508.40, respectively) in the long-chain base, as well as at the Δ2 or Δ3 positions (*m/z* 396.27 or 410.29, respectively) in the *N*-acyl chain (Figure 5a). The sodium adduct EAD showed the co-elution of Cer 18:0;1OH,3OH,4OH/24:0;2OH and Cer 18:0;1OH,3OH,4OH/24:0;3OH at 13.6 min (Figure 5a). In EAD MS/MS analysis, lipid isomers with different C=C positions cannot be quantified using MS2 peaks because uncontaminated MS/MS fragments that distinguish the C=C position cannot be obtained. This is due to the presence of three fragment patterns (radical, H-loss, and H-gain) at ±1.01 Da for each FA moiety. In contrast, the positional isomers of lipids with OH groups can be distinguished by a fragment mass difference of 14.02 Da because the positional difference is at least at the adjacent carbon^37^.

**Figure 5.**
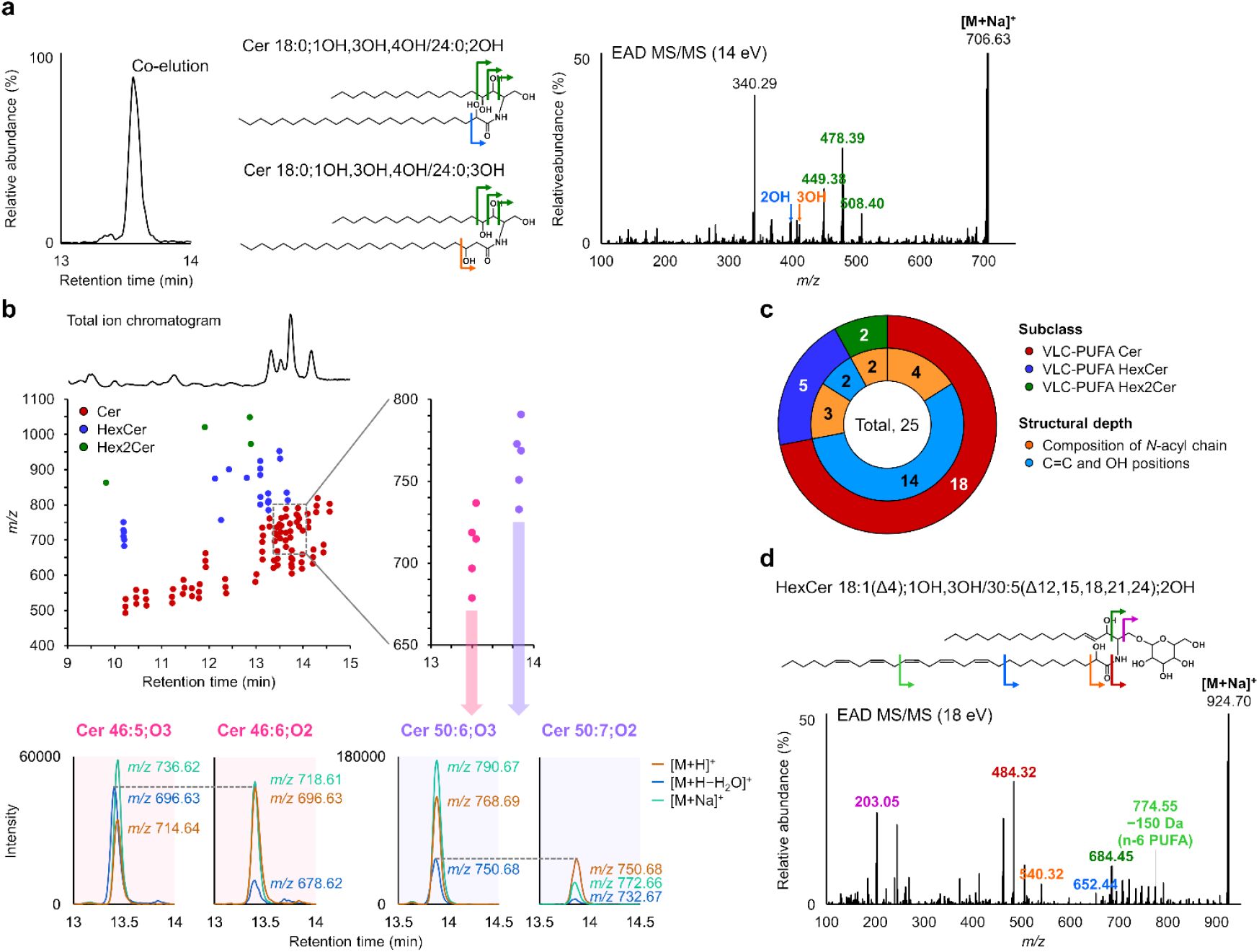
In-depth sphingolipid profiling and structural characterization. (**a**) Structural analysis of Cer 18:0;O3/24:0;O in the mouse feces in the acidic MeOH fraction after SPE enrichment. Co-eluted Cer 18:0;1OH,3OH,4OH/24:0;2OH and Cer 18:0;1OH,3OH,4OH/24:0;3OH (both, 13.6 min) were annotated with specific fragments derived from the OH position of *N*-acyl chains at 14 eV kinetic energy. (**b**) In-depth sphingolipid profiling of mouse testis extract in the acidic MeOH fraction after SPE enrichment. Three adduct forms ([M+H]^+^, [M+H−H_2_O]^+^, and [M+Na]^+^) were used to annotate Cer types. Because dehydrated ions from Cer-HS and protonated ions from Cer-NS (e.g., Cer 46:5;O3 vs. Cer 46:6;O2 and Cer 50:6;O3 vs. Cer 50:7) share identical *m/z* values, several peaks contain contributions from co-eluting isomeric species. (**c**) Number of VLC-PUFA-containing Cer, HexCer, and Hex2Cer molecules detected in sphingolipid-enriched extracts from mouse testis. Details of the detected molecules and their structures are summarized in Table S3. (**d**) Structural analysis of HexCer 18:1;O2/30:5;O in the mouse testis. Omega-6 polyunsaturated hydroxy FA at the *N*-acyl chain was characterized using the 18 eV kinetic energy.

VLC-PUFAs are found in the brain, retina, and testis, where they play a role in nervous system development, photoreceptor function in vision, and spermatogenesis in male reproduction, respectively^17^. VLC-PUFA-containing sphingolipids are synthesized by CerS3 (Cer synthase) and ELOVL2 (FA elongase) in germ cells. AdipoR2 (adiponectin receptor), a key regulator of membrane homeostasis, is essential for maintaining membrane fluidity in male germ cells to ensure proper meiotic progression and male fertility^45^. It also regulates VLC-PUFA synthesis by modulating CerS3 and ELOVL2 expression at both transcriptional and post-transcriptional levels^45^. In the present study, the diversity of VLC-PUFA-containing Cers and hexosylceramides (HexCers) was analyzed in the sphingolipid fraction of mouse testis extract. In mouse testis, dehydrated ions from Cer-HS and protonated ions from Cer-NS, such as Cer 46:5;O3 vs. Cer 46:6;O2 and Cer 50:6;O3 vs. Cer 50:7;O2, cannot be distinguished because they share identical *m/z* values and similar retention times (Figure 5b). Our sodium LC-MS system enables accurate annotation of Cer adduct forms under such co-eluting conditions. In addition, a total of 25 VLC-PUFA sphingolipids, including 18 Cers, five HexCers, and two dihexosylceramides (Hex2Cers), were annotated using EAD MS/MS-based lipidomics (Figure 5c). Among these molecules, the structures of 14 Cers and two HexCer were characterized using the EAD MS/MS system. HexCer 18:1;O2/30:5;O (*m/z* 924.69) analyzed with sodium adduct EAD revealed the OH group at the Δ2 position (*m/z* 540.31) in the *N*-acyl chain (Figure 5d). At a higher kinetic energy of 18 eV, a specific H-gain fragment corresponding to the third C=C position from the methyl terminus (*m/z* 774.55 at the *n*-11 position) was observed, suggesting that the *N*-acyl chain was *n*-6 PUFA (Figure 5d). This fragment pattern, known as vinyl cleavage, has been previously reported in the analysis of *N*-(4-aminomethylphenyl)pyridinium-derivative PUFA using CID MS/MS, which promotes charge-remote fragmentation^46^. These specific fragments suggested the structure as HexCer 18:1(Δ4);1OH,3OH/30:5(Δ12,15,18,21,24);2OH, although the exact structure of the hexose was not determined owing to the lack of glucose- or galactose-specific fragments. A similar C=C and OH group arrangement on the *N*-acyl chain was also observed in Cer 18:1(Δ4);1OH,3OH/30:5(Δ12,15,18,21,24);2OH. Although the structure of Hex2Cer cannot be fully characterized owing to its low abundance, it may share a structure similar to that of VLC-PUFA-containing Cer and HexCer, whose mechanistic roles in glycosphingolipid metabolism remain to be elucidated.

## Conclusions

We developed a sphingolipid profiling method that employs sodium adduct ions alongside proton adduct ions to improve peak annotation using retention time information and to enable detailed structural elucidation via EAD MS/MS. Quantum chemical calculations revealed differences in the dissociation rates, indicating that the dehydration of sodium adduct ions requires a higher activation energy than that of proton adduct ions, even though the resulting product is thermodynamically stable. They also elucidated that the sodium adduct EAD highlights the key fragments by electronic activation and subsequent homolytic allylic cleavage of Cers, enabling the precise structural determination of the C=C and OH positions. Using this approach, we annotated unique Cer species, including AP- and BP-type Cers (Cer-AP and Cer-BP) and *n*-6 VLC-PUFA HexCer-AS, in mouse feces and testis. To better understand the relationship between gut microorganisms and human physiology, region-specific information was obtained through targeted sampling using pH differences across various intestinal regions^47^. Although the diversity of fecal lipids remains unclear, sodium adduct-enhanced EAD may facilitate the understanding of physiological functions, including interactions between microbiota and human metabolic pathways. However, the current system has a limited ability to annotate sugar moieties (e.g., glucose or galactose), presenting a challenge for the comprehensive structural analysis of glycolipids. Recently, mannose-rich glycans and sialic acid linkages in *N*-linked glycopeptides were analyzed using electron-based fragmentation^48,49^. For glycolipid analysis, cross-ring fragmentation of galactosylceramides is observed^33^, providing valuable information for determining the glycosidic linkage positions of long glycans and the acyl chain attachment sites in acylated glycolipids. Our combined approach of integrating quantum chemical calculations with sodium adduct EAD techniques has significant potential for advancing lipidomics. This offers a deeper understanding of adduction sites and their corresponding fragmentation mechanisms, paving the way for more detailed analysis of complex lipid structures.

## Author Contributions

H. Takeda, D. A., and H. Tsugawa designed the study (Conceptualization). D. A. performed the quantum chemical calculations and interpreted the fragmentation pathway (Investigation and Methodology). H. Takeda and M. T. optimized the method, analyzed the lipids, and performed peak annotation and structural analysis (Investigation, Methodology and Data curation). H. Takeda, D. A., and H. Tsugawa wrote the manuscript (H. Takeda and D. A., Writing – original draft; H. Tsugawa, Writing – review & editing) and prepared the figures (Visualization). All the authors approved the final version of the manuscript.

## Data availability

Raw MS data are available in the MB-POST repository (https://repository.massbank.jp/) under the index number MPST000153.

## Supporting Information

The Supporting Information is available free of charge at ACS Publications website.

- Supplemental Figures: Calculated dehydration pathway of protonated ceramides (Figure S1), RPLC/MS system of sphingolipid profiling (Figure S2), and structural analysis of ceramides by sodiated EAD MS/MS (Figure S3).
- Supplemental Tables: Parameters for data analysis using MS-DIAL 5 (Table S1) and deep profiling of ceramides in mouse feces and testis (Tables S2 and S3).

## Acknowledgments

Molecular structure computations were performed at the Research Center for Computational Science in Okazaki, Japan (project: 25-IMS-C065). This study was supported by the Japan Agency for Medical Research and Development (AMED) under Infectious Diseases Research and Infrastructure (JP25wm0325071, H.Tsugawa.), the Japan Science and Technology Agency (JST) Exploratory Research for Advanced Technology (ERATO) (JPMJER2101 to H. Tsugawa.), JST FOREST program (JPMJFR230H to H. Tsugawa.), and the JSPS KAKENHI (23H01996 to D. A. and 24K02011, 24H00043, 24H00392, 24K21269, 25H01425, and 25H01426 to H. Tsugawa.).

## Table of Contents

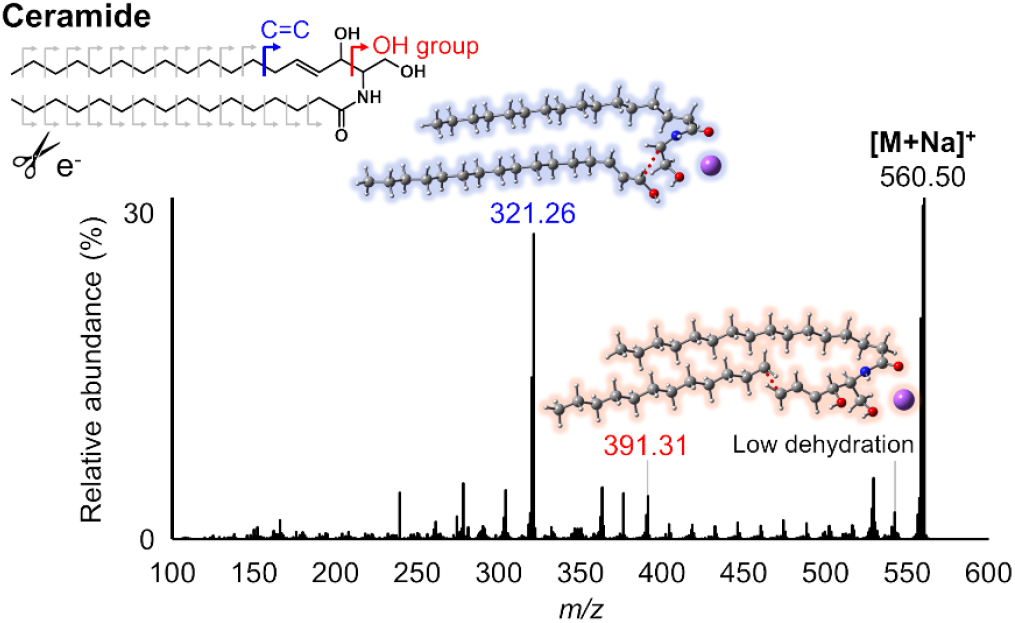

