## Supporting_Information for "Sodium ion-assisted structural lipidomics for sphingolipid profiling"

#### Corresponding Authors

#### Contents

Supplementary Figures 1–3

Supplementary Tables 1–3

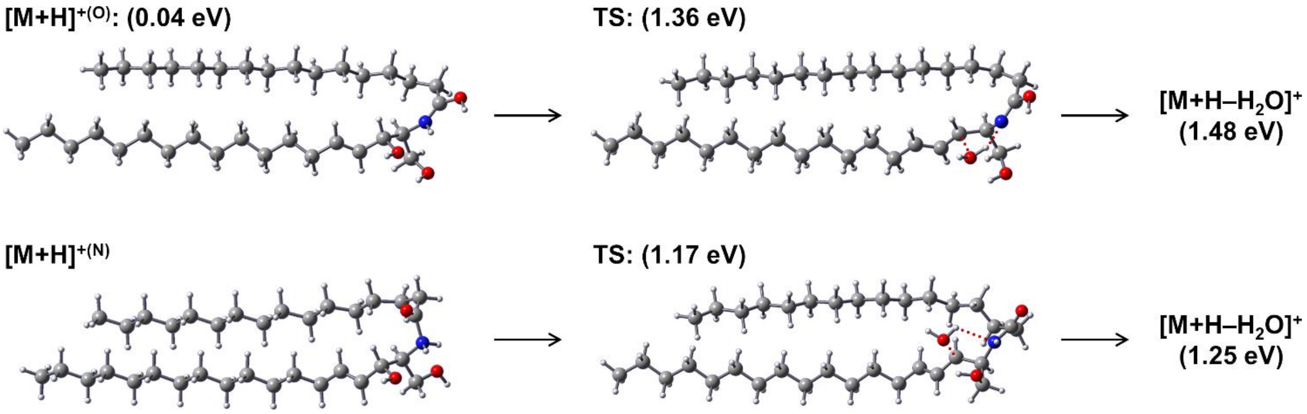


**Figure S1. Calculated dehydration pathway of protonated Cer-NS 18:1(4E);1OH,3OH/16:0.** Carbon, hydrogen, oxygen, nitrogen, and sodium atoms are represented by the gray, white, red, blue, and purple spheres, respectively. Initial energy of O-protomer is referenced to that of N-protomer. Activation energies of transition states and enthalpy changes between reactants and products were calculated for O-protonated and N-protonated forms.

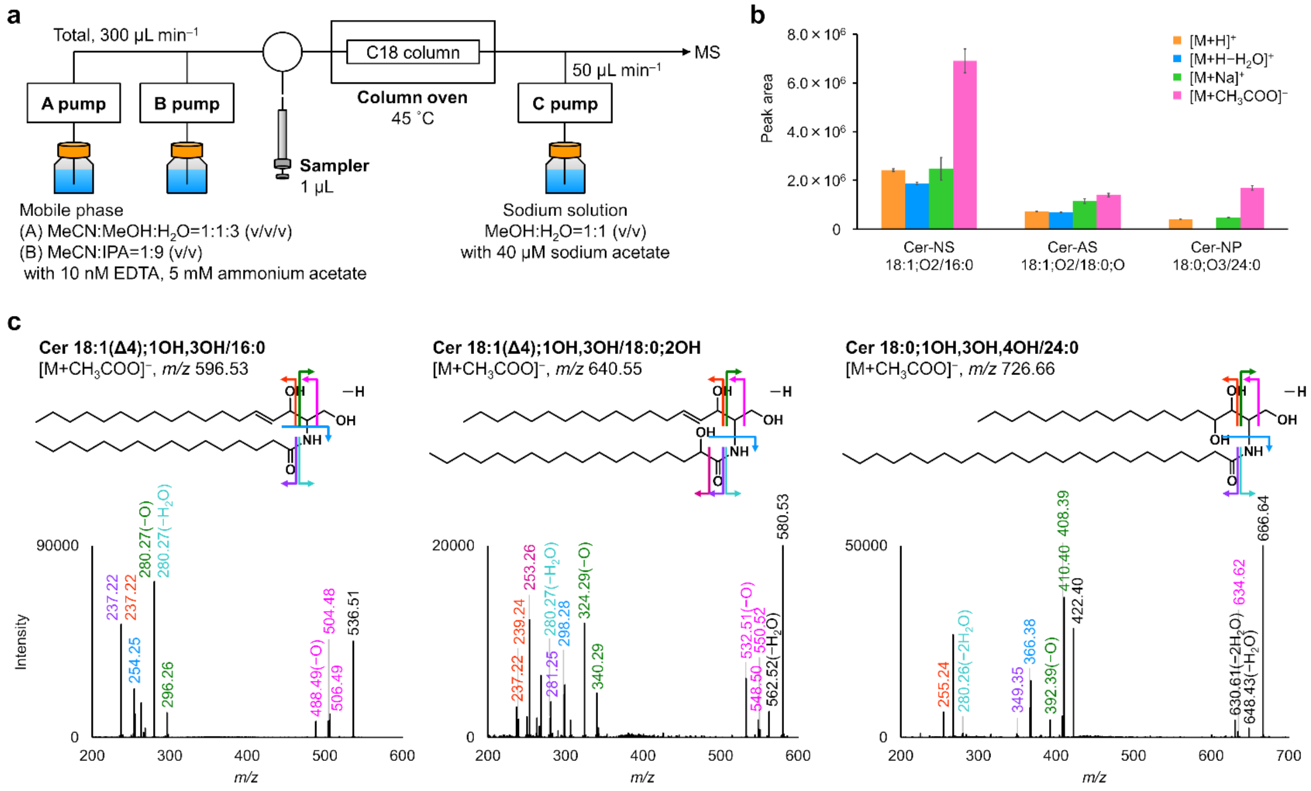


**Figure S2. RPLC/MS system of sphingolipid profiling.** (**a**) Schematic of the LC system for the addition of the sodium solution. (**b**) Comparison of MS1 sensitivities for $[M+H]^+$, $[M+H-H_2O]^+$, $[M+Na]^+$, and $[M+CH_3COO]^-$ ions. (**c**) CID MS/MS spectra of Cers in negative ion mode. In addition to annotating the composition of the long-chain base and the *N*-acyl chain, the OH position on the N-acyl chain was also characterized, based on a previous report[44].

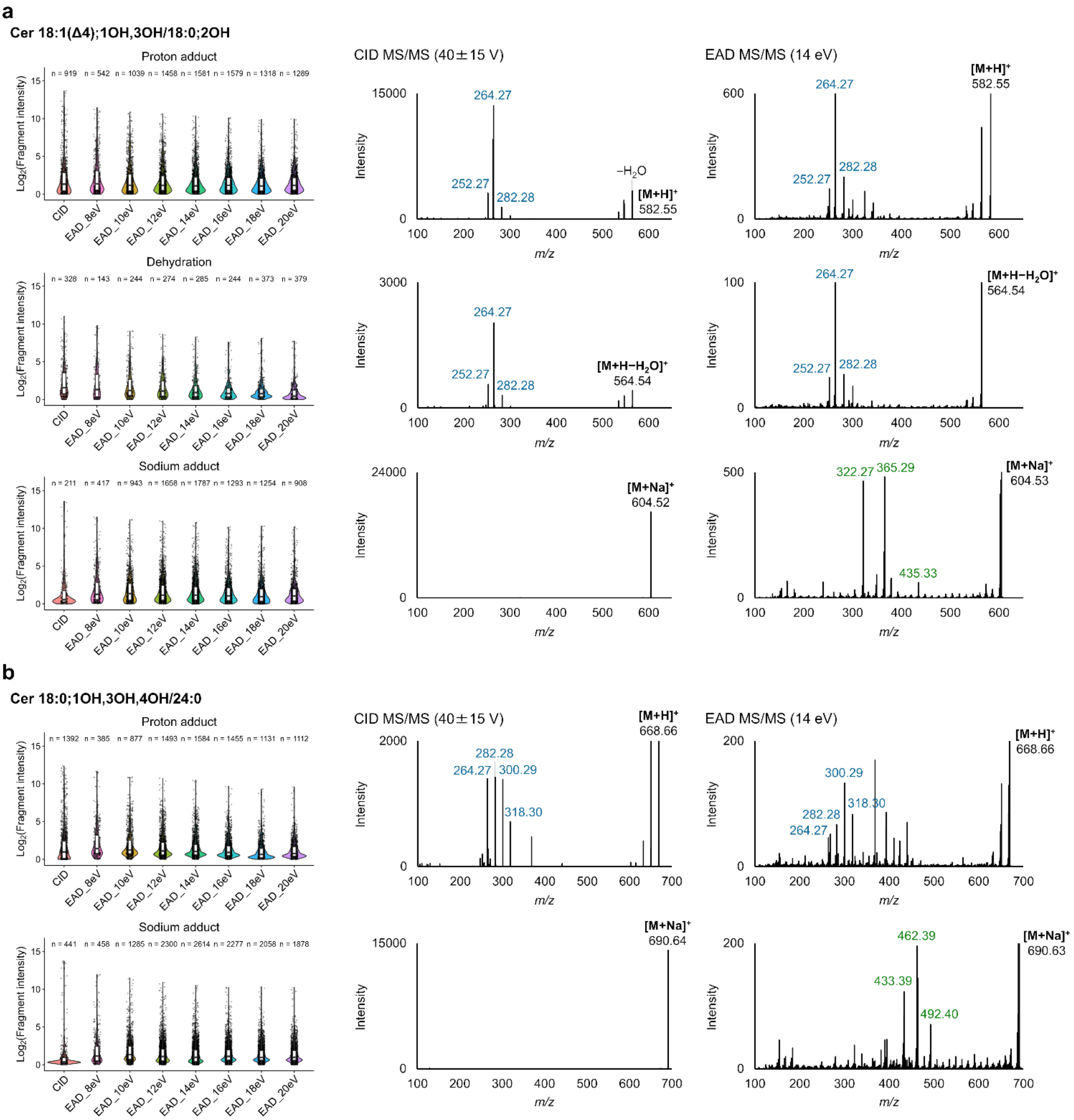


**Figure S3. Advanced structural analysis of three Cer types using sodium adduct EAD MS/MS.** (**a**) Structural analysis of Cer 18:1(Δ4);1OH,3OH/18:0;2OH. The left panel shows the distribution of fragment ion numbers and their intensities for proton and sodium adducts under CID and EAD conditions. Black circles indicate individual fragments with an intensity greater than 1, and the number of such fragments is shown at the top of each plot (n = xx). The right panels show CID MS/MS and EAD MS/MS of the proton adduct, its dehydrated form, and the sodium adduct. Navy blue and green fragments originate from major vibrational and electronic activations, respectively. (**b**) Structural analysis of Cer 18:0;1OH,3OH,4OH/24:0.

### Supplementary Tables

Table S1. Parameters of MS-DIAL software.

Table S2. Structural analysis of ceramides in mouse feces.

Table S3. Structural analysis of ceramides in mouse testis.
